# Biphasic recruitment of TRF2 to DNA damage sites promotes non-sister chromatid homologous recombination repair

**DOI:** 10.1101/305169

**Authors:** Xiangduo Kong, Gladys Mae Saquilabon Cruz, Sally Loyal Trinh, Xu-Dong Zhu, Michael W. Berns, Kyoko Yokomori

**Author notes:** Authors for correspondence: Michael W. Berns, Department of Developmental and Cell Biology, University of California, Irvine. Beckman Laser Institute Irvine, California, 92612, USA; Kyoko Yokomori, Department of Biological Chemistry, School of Medicine, University of California, Irvine, California, USA.

## Abstract

TRF2 binds to telomeric repeats and is critical for telomere integrity. Evidence suggests that it also localizes to non-telomeric DNA damage sites. However, this recruitment appears to be precarious and functionally controversial. We find that TRF2 recruitment to damage sites occurs by a two-step mechanism: the initial rapid recruitment (phase I) and stable and prolonged association with damage sites (phase II). Phase I is poly(ADP-ribose) polymerase (PARP)-dependent and requires the N-terminal basic domain. The phase II recruitment requires the C-terminal MYB/SANT domain and the iDDR region in the hinge domain, which is mediated by the MRE11 complex and is stimulated by hTERT. PARP-dependent recruitment of intrinsically disordered proteins contributes to transient displacement of TRF2 that separates two phases. TRF2 binds to the I-PpoI-induced DNA double-strand break sites, which is enhanced by the presence of complex damage and is dependent on PARP and the MRE11 complex. TRF2 depletion affects non-sister chromatid homologous recombination (HR) repair, but not HR between sister chromatids or non-homologous endjoining pathways. Our results demonstrate a unique recruitment mechanism and function of TRF2 at non-telomeric DNA damage sites.

**Summary Statement:** TRF2 is recruited to DNA double-strand break damage sites by a two-step mechanism and functions in non-sister chromatid homologous recombination repair

## Introduction

TRF2 is an integral component of the telomere shelterin complex that protects telomere integrity (Bilaud, et al., 1997, Feuerhahn, et al., 2015, Okamoto, et al., 2013, van Steensel, et al., 1998). TRF2 recognizes telomere repeat sequence directly through its C-terminal MYB/SANT domain, and protects telomeres by both promoting T-loop formation and inhibiting the DNA damage checkpoint kinase, ATM (de Lange, 2002, Griffith, et al., 1999, Karlseder, et al., 2004, van Steensel, et al., 1998). TRF2 was also shown to be recruited to non-telomeric DNA damage sites and promotes DNA double-strand break (DSB) repair, though its exact role in the process remains controversial. TRF2 recruitment was observed at high-irradiance laser-induced DNA lesions, but not at damage sites induced by low-irradiance ultraviolet radiation or ionizing radiation, despite the presence of DSBs in both cases (Bradshaw, et al., 2005, Huda, et al., 2012, Williams, et al., 2007). TRF2 was linked to HR repair (Mao, et al., 2007), but its phosphorylation by ATM appears to be important for fast repair (suggested to be non-homologous endjoining (NHEJ)) (Huda, et al., 2009). Thus, TRF2 recruitment and function at non-telomeric DNA damage sites remain enigmatic.

PARP1 is a DNA nick sensor activated rapidly and transiently in response to DNA damage (for reviews (Ball and Yokomori, 2011, Beck, et al., 2014, Daniels, et al., 2015, Kalisch, et al., 2012)). There are multiple PARP family members, but PARP1 plays a major role in PAR response at damage sites (Cruz, et al., 2015, Kong, et al., 2011). Activated PARP1 uses NAD^+^ as a substrate to ADP-ribosylate multiple target proteins, including itself. Although PARP1 was initially thought to specifically facilitate base excision repair/single-strand break repair, recent studies reveal its role in multiple DNA repair pathways, including DSB repair (Beck, et al., 2014). Furthermore, PAR modification at DNA damage sites is critical for the recruitment of chromatin modifying enzymes that promote DNA repair (Ahel, et al., 2009, Ayrapetov, et al., 2014, Ball and Yokomori, 2011, Chou, et al., 2010, Gottschalk, et al., 2009, Izhar, et al., 2015, Khoury-Haddad, et al., 2014, Larsen, et al., 2010, Polo, et al., 2010, Smeenk, et al., 2010, Sun, et al., 2009). Thus, PARP1 is not only a sensor of DNA damage, but also a regulator of damage site chromatin environment and multiple DNA repair pathways in higher eukaryotes.

## Results and Discussion

### TRF2 recruitment to damage sites is determined by the degree of PARP activation

Previously, we found that TRF2 is rapidly recruited to higher input-power laser damage sites that contained complex DNA damage in a PARP-dependent manner (Cruz, et al., 2015). In contrast, lower input-power laser irradiation that induced relatively simple strand-breaks and no significant PAR response failed to recruit TRF2 (for laser power measurements, see **Materials and Methods**). Importantly, stimulation of PARylation at lower input-power laser damage sites by a PARG inhibitor promotes TRF2 recruitment (Fig. 1A). Therefore, the level of PARP activation, rather than the nature of damage per se, is the deciding factor for TRF2 recruitment to non-telomeric DNA damage sites.

**Figure 1.**
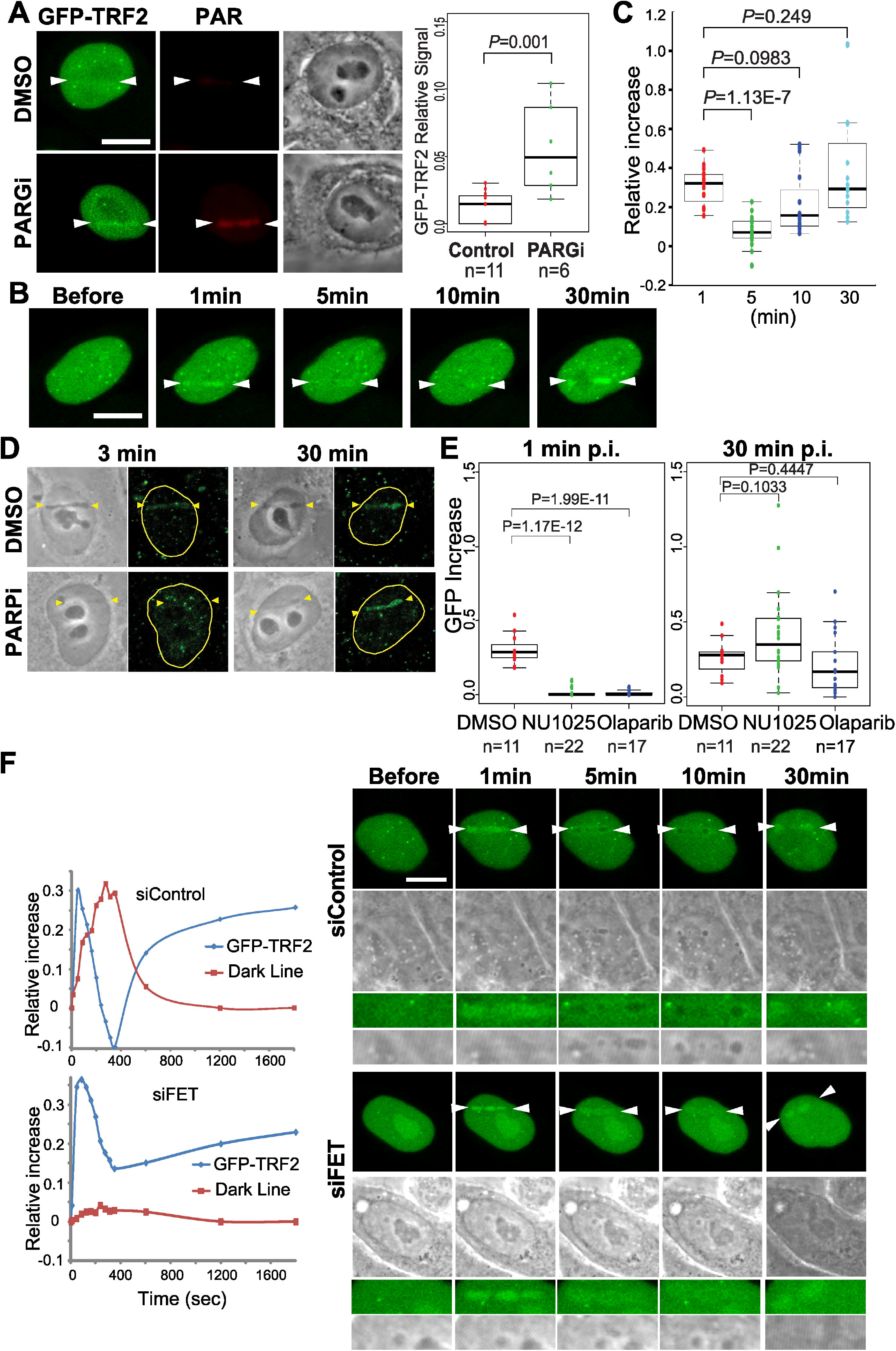
Biphasic TRF2 recruitment to non-telomeric damage sites. **A.** PAR stimulation by PARG depletion promotes GFP-TRF2 accumulation at low input-power damage sites. Scale bar is 10 μm in all figures. Quantification of the relative increase of GFP signals at damage sites is shown. **B.** Time course analysis of GFP-TRF2 recruitment to laser-induced DNA damage sites. **C.** Quantification of GFP signals at damage sites as in (A). N=16 **D.** Detection of the endogenous TRF2 at damage sites. PARP inhibition suppresses phase I, but has no effect on phase II, TRF2 recruitment. **E.** The effects of PARP inhibitors (NU1025 and olaparib) on immediate (phase I) and late (phase II) GFP-TRF2 recruitment. **F.** The effect of IDP depletion on dispersion of TRF2 at damage sites. Right: Time course analysis of signal intensity changes of GFP-TRF2 (blue) and dark line (red) in cells transfected with control siRNA (siControl) or FET siRNAs (siFET).

### Biphasic recruitment of TRF2 to damage sites

We and others observed rapid and transient TRF2 recruitment to damage sites within the first 5 min post irradiation (p.i.) (Bradshaw, et al., 2005, Cruz, et al., 2015) (Fig. 1). Upon inspection of later time points (20-30 min), however, we found that TRF2 re-appears at damage sites (Fig. 1B and C). Similar recruitment patterns were observed with both the ectopically expressed and endogenous TRF2 (Fig. 1B and D, respectively). The initial recruitment of GFP-TRF2 peaks at ~1-3 min p.i. (termed “phase I”), which decreases once but returns peaking at ~30 min to 1 hr (“phase II”) (Fig. 1A-C). This phase II recruitment persists for at least 2 hr (data not shown). Interestingly, the PARP inhibitors NU1025 and olaparib completely suppressed phase I, but not phase II, recruitment of both endogenous and recombinant TRF2 (Fig. 1D and E), suggesting that two phases of TRF2 recruitment are mediated by different mechanisms.

### IDPs compete with TRF2 for PARylated DNA lesions

We found that transient GFP-TRF2 displacement is inversely correlated with the appearance of a prominent dark line at damaged lesions readily visible using bright field microscope imaging (Fig. 1B and F). Close examination at damage sites revealed that GFP signals not only decrease, but are often transiently pushed to the periphery of the damage sites, suggesting that it may be displaced by the constituents of the dark line (Fig. 1B and F). Intrinsically disordered proteins (IDPs), FUS, EWS and TAF15 (FET), were shown to accumulate at damage sites in a PAR-dependent manner and are the major components of the dark line (Altmeyer, et al., 2015). We found that depletion of FET by siRNA resulted in a more even distribution of the GFP-TRF2 signal at damaged lesions, correlating with disappearance of the dark line (Figs. 1F and S1A). The results reveal that although both TRF2 and IDPs are recruited to damage sites in a PAR-dependent manner, there is a distinct order of appearance and competition between them, which separates phases I and II.

### Phase I and II recruitments are mediated by distinct domains

TRF2 protein domains have been characterized extensively in the context of telomeres. The C-terminal MYB/SANT DNA binding domain of TRF2 specifically recognizes and binds telomere DNA whereas the N-terminal basic domain is not required for telomere targeting (Fig. 2A) (Karlseder, et al., 1999, Okamoto, et al., 2013). We found that the phase I recruitment is abolished by deletion of the N-terminal basic domain similarly to that previously reported (Bradshaw, et al., 2005) (Fig. 2B and C). In contrast, phase II recruitment was significantly inhibited by deletion of the C-terminal MYB/SANT domain. Deletion of both N- and C-terminal domains abolished both phases of damage site recruitment. Thus, distinct domains play critical roles in rapid and transient phase I and slow and stable phase II. Unlike the previous report (Bradshaw, et al., 2005), we found that deletion of the MYB/SANT domain partially reduced phase I recruitment, suggesting that the phase I recruitment is further stabilized by DNA binding of the MYB/SANT domain (Fig. 2C).

**Figure 2.**
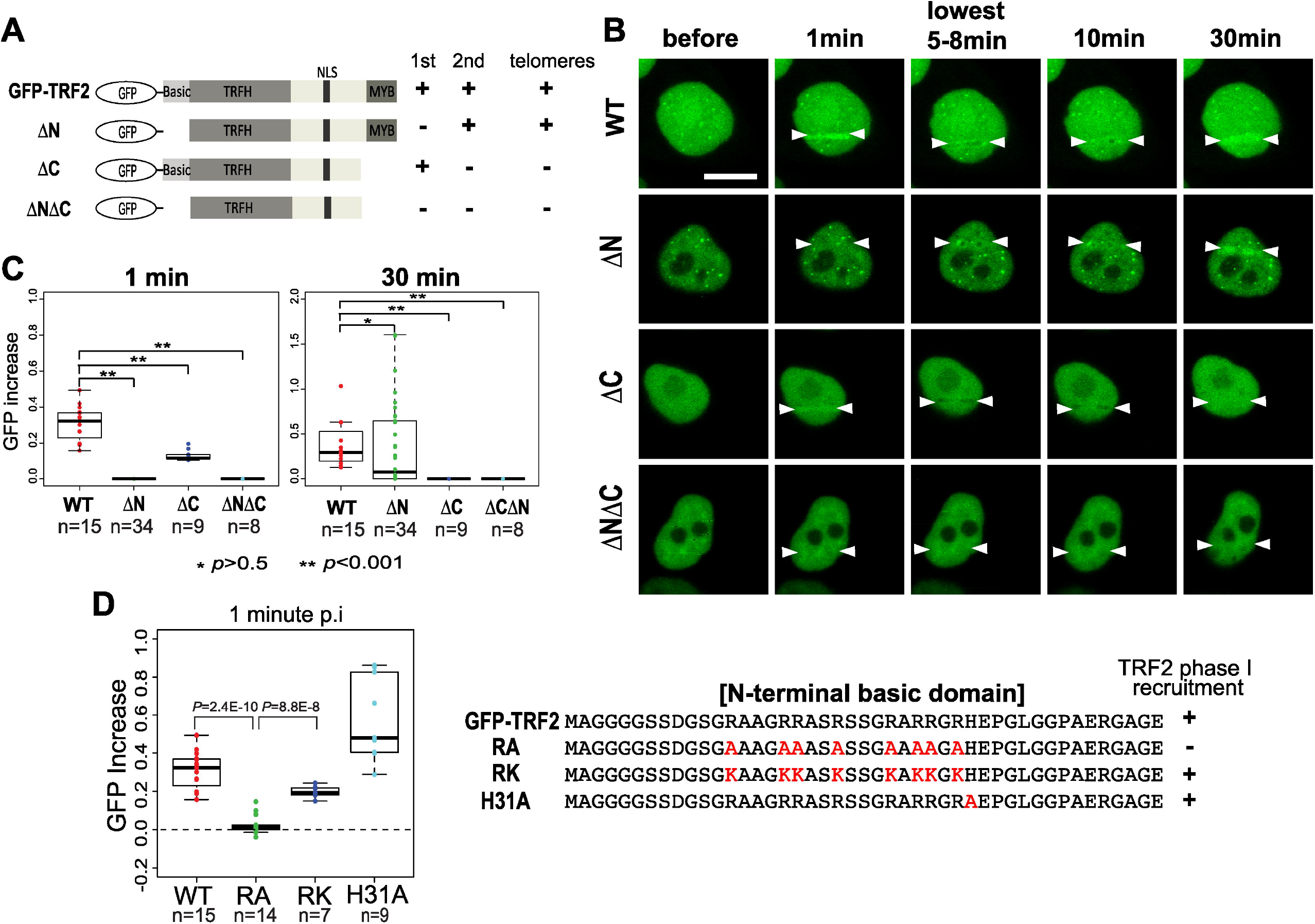
Distinct domain requirement for phase I and II recruitment. **A.** Schematic diagrams of TRF2 deletion mutants. **B.** Time course analysis of damage site localization of WT and deletion mutants. **C.** Quantification of the mutant GFP signals at damage sites at 1 min (phase I) and 30 min (phase II) post damage induction. **D.** The effects of the N-terminal amino acid substitutions on damage on phase I recruitment. Arginine to alanine mutations (RA), arginine to lysine substitution (RK), the HJ binding mutation (H31A) were tested.

### Positive charge of the basic domain is required for phase I recruitment

The N-terminal basic domain harbors multiple arginine residues, which may interact with negatively charged PAR residues clustered at damage sites. Indeed, arginine to alanine mutations completely abolished the phase I recruitment (Fig. 2D, “RA”). In contrast, this recruitment is sustained albeit weaker by arginine to lysine mutations that preserve the positive charge (Fig. 2D, “RK”). The basic domain was also shown to bind to the holiday junction (HJ) (Fouché, et al., 2006). However, HJ binding-defective mutation (H31A) (Poulet, et al., 2009) showed no inhibitory effect on TRF2 recruitment to damage sites (Fig. 2D). Thus, the positive charge is essential for the PARP-dependent TRF2 recruitment to non-telomeric DNA damage sites.

### hTERT contributes to MYB/SANT-dependent phase II recruitment

Human telomerase reverse transcriptase (hTERT) is responsible for addition of telomere sequences. Several studies indicated that it can also polymerize DNA at non-telomeric DNA ends de novo and has a distinct role in DNA damage response (DDR) and repair (Flint, et al., 1994, Gao, et al., 2008, Majerská, et al., 2011, Masutomi, et al., 2005, Morin, 1991, Ribeyre and Shore, 2013). Since the MYB/SANT domain, which is responsible for telomere repeat recognition, is pertinent to phase II recruitment, we tested the possible contribution of hTERT in the TRF2 association at DSB sites. HeLa cells, which express hTERT, were treated with siRNA specific for hTERT (Figs. 3A and S2A). We found that the phase II recruitment of TRF2 was partially inhibited by this treatment (Fig. 3A). TRF2, however, can also be recruited to damage sites even in telomerase-negative ALT cells albeit at a lower level (e.g., U2OS cells) (Fig. S2B). Thus, the results indicate that hTERT contributes to, but is not essential for, the MYB/SANT domain-dependent TRF2 recruitment to damage sites.

**Figure 3.**
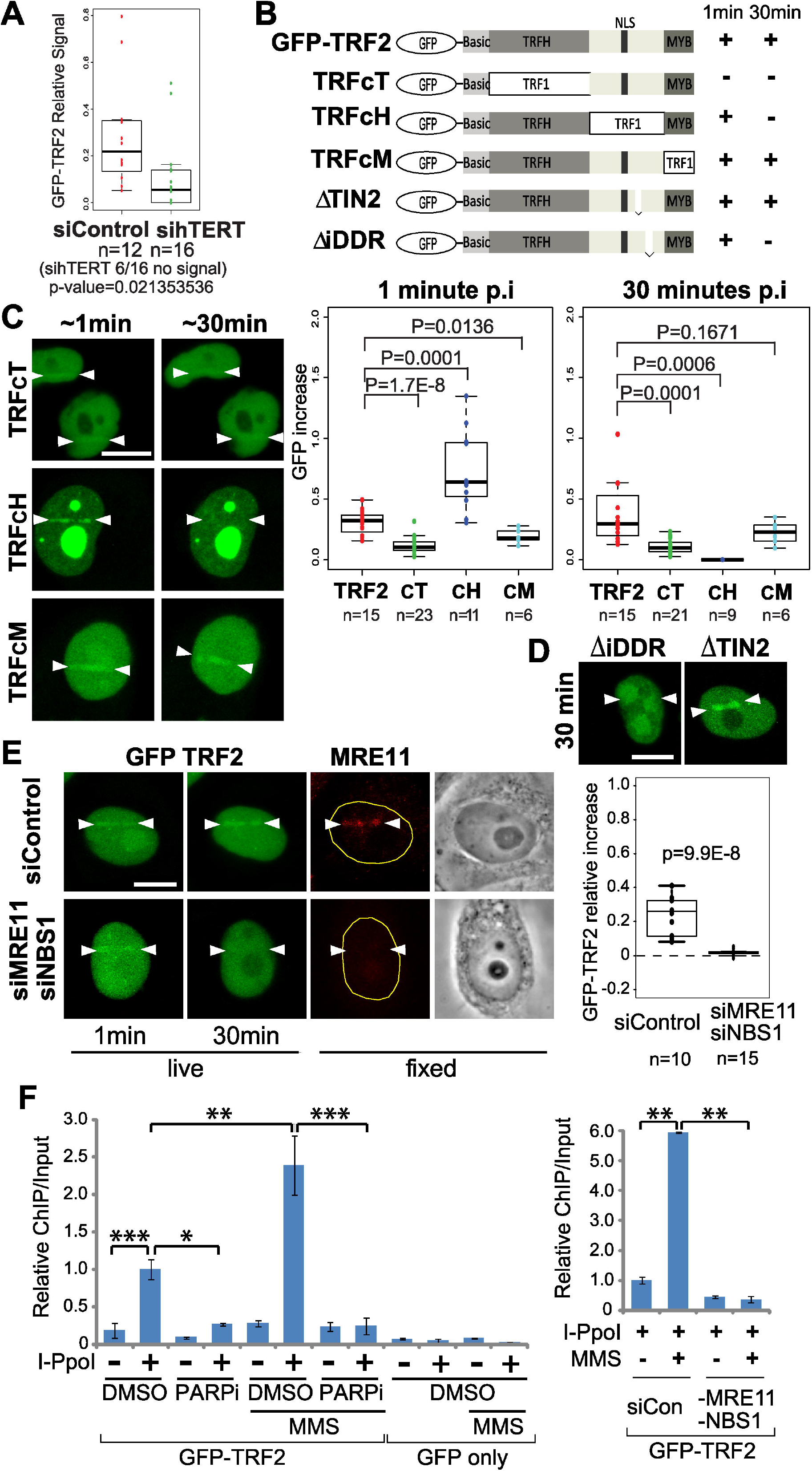
Phase II recruitment is affected by hTERT and is dependent on the iDDR region in the hinge domain of TRF2. **A.** hTERT depletion by siRNA inhibits TRF2 phase II recruitment. **B.** Schematic diagrams of TRF2 mutants (Okamoto, et al., 2013). **C.** Representative cell images of the recruitment of chimeric mutants to damage sites at ~1 min (phase I) and 30 min (phase II) after damage induction. Quantification of the GFP-TRF2 signal increase at damage sites is shown. **D.** Comparison of iDDR and TIN2 deletion mutants at 30 min after damage induction. **E.** The effect of MRE11 and NBS1 siRNA depletion on phase I and II recruitment of GFP-TRF2 was examined comparing to siControl. Cells were fixed and stained with anti-MRE11 antibody to confirm the depletion. Quantification of the GFP-TRF2 signal increase at damage sites in control or Mre11/NBS1 siRNA-treated cells is shown. **F.** ChIP-qPCR analysis of GFP-TRF2 binding at I-PpoI cut sites. TRF2 binding was examined in the absence or presence of I-PpoI, and with and without MMS as indicated. Cells were further treated with DMSO or PARP inhibitor (PARPi) (left panel). GFP only was used as a negative control. Alternatively, cells were transfected with control (siCON) or MRE11/NBS1 siRNA in the presence of I-PpoI with or without MMS (right panel). * p < 0.01, ** p < 0.001, *** p < 0.0001.

### Additional domain requirements for phase I and II recruitments

Chimeric mutants between TRF1 and TRF2 have provided important insight into the unique TRF2 function in telomere protection (Okamoto, et al., 2013). The N-terminal basic domain is unique to TRF2 (TRF1 has the acidic domain), indicating that phase I recruitment is specific to TRF2. Similarly, we found that the TRF2 TRFH domain required for dimerization (<30% homology with TRF1) is essential for both phase I and II recruitment (Fig. 3B and C; TRFcT). Interestingly, the TRFcH mutant replacing the TRF2 hinge domain with that for TRF1 exhibited intact phase I recruitment, but failed for phase II, indicating that the hinge domain (11% homology to TRF1) is uniquely required for the latter recruitment. In contrast, the TRF2 MYB/SANT domain was found to be interchangeable with that for TRF1 for phase II recruitment (both recognize telomere repeats) (Fig. 3B and C; TRFcM).

The hinge domain contains binding sites critical for several different factors (Chen, et al., 2008, Okamoto, et al., 2013). Deletion of the TIN2 binding region (a.a. 352–367) (ΔTIN2) critical for TRF2 incorporation into the shelterin complex (Kim, et al., 2004, Liu, et al., 2004, Ye, et al., 2004) had no effect on damage site recruitment, suggesting that TRF2 recruitment to damage sites is independent of the shelterin complex (Fig. 3D). The hinge domain also contains a region critical for suppression of DNA damage response and telomere maintenance (termed “inhibitor of DDR (iDDR)” (a.a. 406-432)) (Okamoto, et al., 2013). Deletion of this domain (ΔiDDR) recapitulated the phenotype of the TRFcH mutant, inhibiting only the phase II recruitment (Fig. 3D). The iDDR region was shown to be necessary and sufficient for the TRF2 interaction with the MRE11 complex (Okamoto, et al., 2013). We found that siRNA depletion of MRE11 and NBS1, the two components of the complex, effectively reduced the phase II recruitment (Figs. 3E and S1B). The results indicate that the MYB/SANT domain-dependent phase II requires the interaction of the iDDR region with the MRE11 complex.

### Both phase I and II mechanisms are required for TRF2 accumulation at the I-PpoI-induced DSB sites

We found TRF2 binding/localization to the actual DSB sites induced by I-PpoI endonuclease in the ribosomal DNA region (Berkovich, et al., 2008) by chromatin immunoprecipitation (ChIP)-qPCR (Fig. 3F). Intriguingly, this binding was further enhanced by methyl methanesulfonate (MMS) treatment, which induces complex damage and stronger PARP activation than simple DSBs. Importantly, TRF2 accumulation at I-PpoI target sites both in the presence and absence of MMS treatment was effectively suppressed by PARP inhibitor, confirming that TRF2 binding to DSB sites is PARP-dependent (Fig. 3F). Interestingly, depletion of the MRE11 complex also abolished TRF2 binding to DSB sites. The results indicate that the strong PARP activation is the key determinant for the efficient TRF2 recruitment to DSB sites and that both phases I and II recruitment mechanisms are important for the stable binding of TRF2 to DSB sites.

### TRF2 facilitates intra-chromosomal HR repair

To determine the significance of TRF2 recruitment to damage sites, the effects of TRF2 depletion were examined using the cell-based assays for different pathways of DSB repair (Hu and Parvin, 2014). TRF2 was depleted for 48 hr before repair assays in order to minimize telomere erosion, which was typically assayed 4-7 days after depletion (Okamoto, et al., 2013, Rai, et al., 2016) (Fig. S3). TRF2 was implicated previously in both HR repair (Mao, et al., 2007) and fast repair (suggestive of NHEJ) (Huda, et al., 2009). Interestingly, we found that TRF2 depletion reduced the efficiency of HR in the I-SceI-dependent HR assay, but not in the sister chromatid exchange (SCE) assay (Fig. 4A and B). The I-SceI assay selectively captures intra-chromatid or unequal sister chromatid HR whereas the SCE assay specifically detects sister chromatid conversion (Potts and Yu, 2005). Furthermore, TRF2 depletion had no significant effect on either classical or alternative NHEJ repair (Fig. 4B). Thus, the results indicate that TRF2 plays a specific role in non-sister chromatid HR. Under our condition, the effect of TRF2 depletion on I-SceI HR was comparable to that of BRCA1 depletion and can be complemented by the wild type TRF2 (Figs. 4A and S1C). BRCA1 is a known promoter of HR (Moynahan, et al., 1999). While either the N- or C-terminal deletion mutant, which primarily affects phase I or phase II, respectively, exhibited variable results, the cT mutant that hinders both phases clearly failed to complement, indicating that both phases are critical for its optimal repair activity (Figs. 2C, 3C and 4). This is consistent with the requirement of both phases for stable TRF2 binding at DSB sites (Fig. 3F).

**Figure 4.**
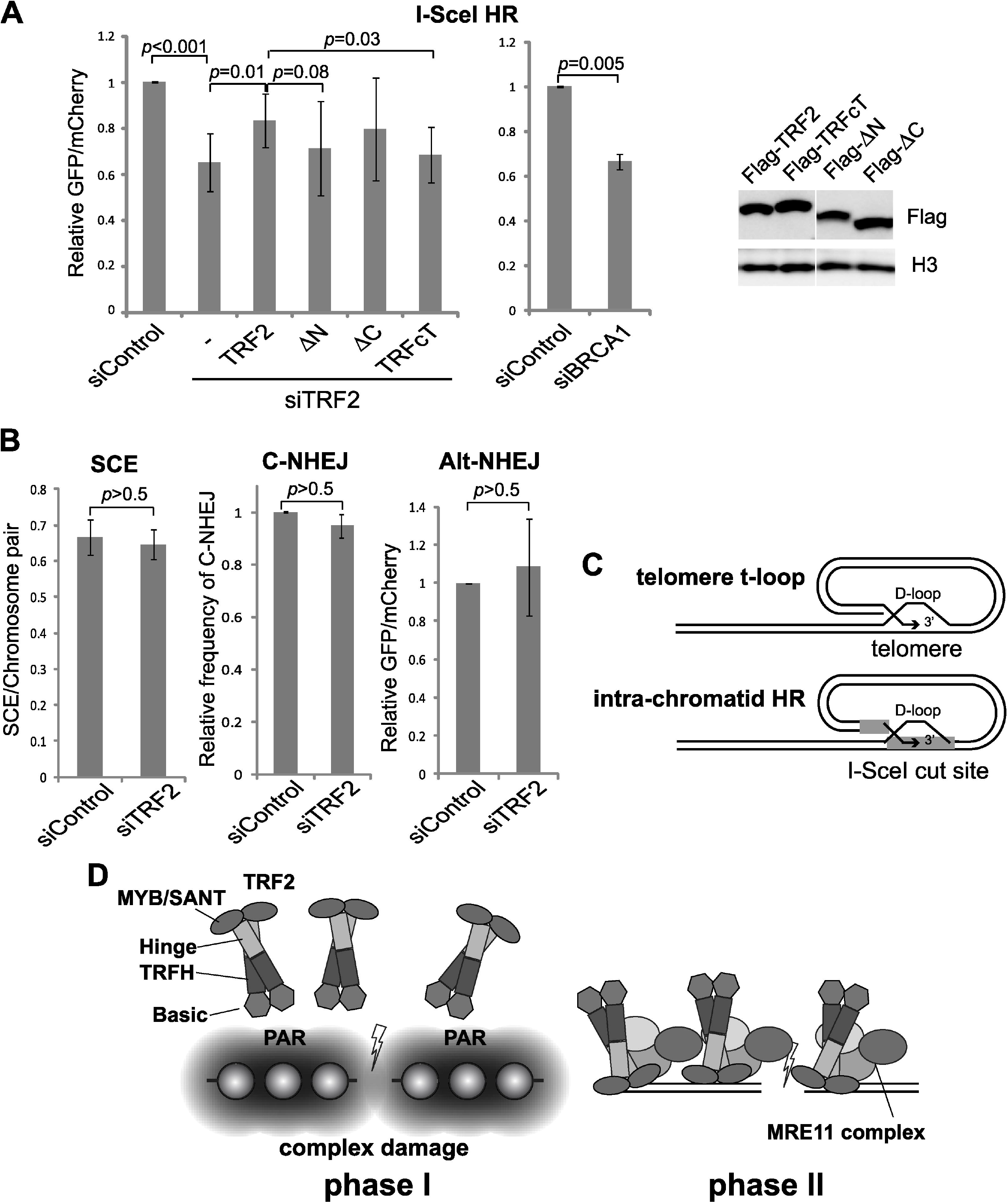
TRF2 specifically promotes non-sister chromatid HR repair. **A.** The effect of TRF2 depletion on DSB repair using the I-SceI HR system. Complementation analysis of TRF2-depleted cells was performed using the wild type and mutants. BRCA1 depletion was used as a control. Comparable expression of the recombinant TRF2 proteins were confirmed by western blot analysis. Histone H3 serves as a loading control. **B.** The effect of TRF2 depletion on different DSB repair pathways was examined using SCE, NHEJ, and alt-NHEJ assays. **C.** Similarity between strand invasion in D-loop formation at telomeres and at DSB sites by TRF2. **D.** Biphasic mechanism of TRF2 recruitment to damage sites. Phase I involves PARP-dependent recruitment through the basic domain. Phase II is mediated by the MYB/SANT domain, which is also dependent on the iDDR region and the Mre11 complex.

The selective involvement of TRF2 in non-sister chromatid HR is an interesting contrast to cohesin, which only promotes sister chromatid HR but not other types of HR (Kong, et al., 2014, Potts, et al., 2006). These can be explained by their mechanisms of actions. Cohesin promotes sister chromatid cohesion, therefore promoting pairing of damaged and undamaged sister chromatids for HR. TRF2 promotes loop formation and strand invasion not only at telomere T-loop, but also with non-telomeric templates in vitro (Amiard, et al., 2007, Doksani, et al., 2013, Griffith, et al., 1999, Stansel, et al., 2001) (Fig. 4C). Furthermore, TRF2 was shown to inhibit Rad51-mediated D-loop formation with a telomeric, but not non-telomeric, template in vitro (Bower and Griffith, 2014). Thus, the cell may hijack the ability of TRF2 by clustering the protein to DSB sites to promote intra-chromatid HR, which may be particularly important in the context of complex damage that robustly activates the PARP response.

Our results demonstrate that the initial phase I recruitment by PARP may be critical to bring TRF2 to the proximity of non-telomeric damage sites, which is then further stabilized by the Mre11 complex interaction and DNA binding through the MYB/SANT domain (Fig. 4D). PAR-dependent accumulation of IDPs may compete with phase I recruitment and trigger the second mode of TRF2 association with DNA lesions. Thus, PARP initiates the cascade of dynamic recruitment of factors, some of which compete with each other, to fine-tune repair pathway choice. Interestingly, a recent study showed that BLM is also recruited to damage sites in a biphasic fashion, initially by the ATM signaling, and subsequently by the MRE11 complex (Tripathi, et al., 2018), suggests that this type of two-step mechanism may be commonly used to ensure versatility and specificity of the factor recruitment.

### Conclusion

Our results demonstrate PARP- and the MRE11 complex-dependent recruitment of TRF2 to DSB sites, reconciling previous controversies and revealing uniquely regulated non-telomeric and shelterin-independent function of TRF2 in non-sister chromatid HR repair.

## Materials and Methods

### Cell lines and synchronization

HeLa and U2OS cells (ATCC) were grown in Dulbecco’s modified Eagle’s medium (DMEM; Gibco) supplemented with L-Glutamine, 10% fetal bovine serum (FBS) and antibiotics. HeLa DR-GFP (for HR assay) (Hu and Parvin, 2014, Pierce, et al., 2001) and EJ2-GFP (for Alt-NHEJ assay) (Bennardo, et al., 2008) stable cells were grown in DMEM with high glucose (4500 mg/l), supplemented with 10% (v/v) FBS, 1% Pen/Strep, 1% GlutaMAX and 1.5 μg/ml Puromycin. 293/HW1 cells (for c-NHEJ assay) (Zhuang, et al., 2009) were grown in DMEM (high glucose), 1% Pen/Strep, 1% Sodium pyruvate, 1% GlutaMAX, 10% FBS, and 2 μg/ml Puromycin. All the cell lines were mycoplasma-negative.

### TRF2 mutants

The expression plasmids containing GFP-TRF2 full-length, deletion or TRF1 chimeric mutants were kindly provided by Dr. Eros Lazzerini Denchi (The Scripps Research Institute, La Jolla, California) (Okamoto, et al., 2013). These cDNAs were also re-cloned into a pIRES vector containing N-terminal FLAG epitope. Those include TRF2ΔN (45a.a-500a.a), TRF2ΔC (1a.a -454a.a), TRF2ΔNΔC (45a.a-454a.a), ΔTIN2, and TRFcT (Okamoto, et al., 2013). RA and RK mutants were described previously (Mitchell, et al., 2009).

### Antibodies

Mouse monoclonal antibodies specific for PAR polymers (BML-SA216–0100, Enzo Life Sciences, Inc., 1:500 dilution for immunofluorescent staining (IF)), TRF2 (NB100– 56506, Novus Biologicals, 1:500 dilution for IF), MRE11 (GTX70212, GeneTex, Inc. 1:500 dilution for IF, 1:1000 for WB), GFP (632592 TAKARA BIO. 1:500 dilution for chromatin immunoprecipitation (ChIP)), Actin (A4700, Sigma, 1:1000 dilution for western blot (WB)), Flag (F3165, Sigma, 1:1000 dilution for WB), and BRCA1 (GTX70111, GeneTex, Inc, 1:1000 dilution for WB) as well as rabbit polyclonal antibodies specific for γH2AX (GTX628789, GeneTex, Inc, 1:500 dilution for IF), EWSR1 (GTX114069, GeneTex, Inc, 1:1000 dilution for WB), TAF15 (GTX103116, GeneTex, Inc, 1:1000 dilution for WB), H3 (14-411, Upstate Bio. 1:1000 dilution for WB) and Rad21 (Kong, et al., 2014) were used.

### Immunofluorescent staining

At different time points after damage induction, cells were fixed in 4% paraformaldehyde (15 min at 4°C), permeabilized in 0.5% Triton X-100 for five min (4°C), and stained with antibodies. The staining procedure was described previously (Kim, et al., 2002). Fluorescent images were captured through a 100× Ph3 UPlanFI oil objective lens (NA, 1.3; Olympus) on a model IX81 Olympus microscope with a charge-coupled device camera.

### Western blot

Protein samples were subjected to SDS-PAGE and then transferred to nitrocellulose membranes as described previously (Schmiesing, et al., 1998). The membranes were blocked with Pierce Protein-Free T20 (PBS) Blocking Buffer (Thermo Fisher Scientific).

The primary antibody was incubated in 3% BSA–0.05% Tween 20 in PBS for 1hr at room temperature or overnight at 4C°, followed by three washes in PBS–0.05% Tween 20. The secondary antibody conjugated with HRP (Promega) was incubated in 3% BSA–0.05% Tween 20 in PBS for 1 hr at room temperature. The filter was then washed three times in PBS–0.05% Tween 20 and developed with SuperSignal West Pico Chemiluminescent Substrate (Thermo Fisher Scientific). Images were acquired using the Image Analyzer (LAS-4000, Fujifilm Co.) and analyzed using Quantity One.

### Laser damage induction and cell imaging

Near-infrared (NIR) femtosecond laser irradiation was carried out using a Zeiss LSM 510 META multiphoton-equipped (3.0-W 170-fs coherent tunable Chameleon Ultra-NIR laser) confocal microscope. The Chameleon NIR beam was tuned to 780 nm, where the software bleach function was used to target linear tracts inside the cell nuclei for exposure to single laser scans (12.8 μs pixel dwell time) through the 100× objective lens (1.3 NA Zeiss Plan APO) (Cruz, et al., 2015). The Peak irradiance at the focal point for the higher input laser power is 5.27×10^10^ W/cm^2^, and for the lower input laser power is 3.24×10^10^ W/cm^2^ (Cruz, et al., 2015). Recruitment of GFP-TRF2 wt and mutant proteins to damage sites was analyzed by live-cell confocal scanning with the 488-nm CW argon laser on the same Zeiss META platform. Fluorescent measurement of the recruitment of GFP-tagged proteins to damage sites was performed by live-cell confocal scanning with the 488-nm CW argon laser on the Zeiss LSM 510 META platform. The signals were measured with the LSM510 software (version 4.0). The data were collected in three separate experiments. P values were calculated by two-tailed t-test using Excel software. Boxplot were created with R program.

### siRNA depletion

HeLa cells were transfected twice 24 h apart with siRNAs at a final concentration of 5 nM using HiPerFect transfection reagent according to the manufacturer’s instructions (Qiagen). siRNAs directed against hTERT (5-TTTCATCAGCAAGTTTGGA-3) (Masutomi, et al., 2005), MRE11 (5-GCTAATGACTCTGATGATA-3) (Myers and Cortez, 2006), NBS1 (5’-GAAGAAACGTGAACTCAAG-3’) (Myers and Cortez, 2006), hTRF2 (SI00742630, Qiagen), BRCA1 (5’-GCTCCTCTCACTCTTCAGT-3’) (Hu and Parvin, 2014), FET (FUS (s5402), EWSR1 (s4886), and TAF15 (s15656), Ambion/LifeTechnologies) (Altmeyer, et al., 2015), and a negative-control siRNA (Qiagen) were used. Cells were harvested for western blot analyses or were subjected to laser microirradiation, approximately 48 h after the final transfection.

### Inhibitor treatment

20 μM PARP inhibitor olaparib (Apexbio Technology) or 1 μM PARG inhibitor (PARGi) DEA ((6,9-diamino-2-ethoxyacridine lactate monohydrate) (Trevigen)) was added to the cell culture one hour prior to damage induction. DMSO only was added to control cells.

### DSB induction by I-PpoI endonuclease and ChIP-qPCR analysis

The experimental procedure was previously described (Kong, et al., 2014). Briefly, HeLa cells were transfected with the GFP-TRF2 (or GFP only) with or without ER-I-PpoI expression plasmid. 24 hrs later, 4-hydroxy-tamoxifen (4-OHT; Sigma) was added to a final concentration of 1 *μ*M for 10 hr to induce DNA damage. For MMS-treated cells, 3mM MMS was added 1hr before harvest. DMSO or PARP inhibitor was added 1hr before MMS treatment. For MRE11/NBS1 depletion experiment, two rounds of control or MRE11/NBS1siRNA transfection were performed 48 and 24 hrs before GFP-TRF2/ ER-I-PpoI transfection. ChIP was performed, using GFP antibody, as described previously (Berkovich, et al., 2008, Kong, et al., 2014). The data represent the means ± the s.d. of two separate experiments. Four samples (absence or presence of I-PpoI, and with and without MMS) were repeated the third time, and results with same trend were obtained. P values were calculated by two-tailed t-test.

### DSB repair assays

Homologous Recombination and Alt-NHEJ assays were performed as described previously in HeLa cells (Bennardo, et al., 2008, Hu and Parvin, 2014) with modification. Briefly, on day 1, the appropriate cell lines were seeded in 24-well plates. The next day, cells, 50% confluent, were transfected with siTRF2 or siControl. On day 3, cells were re-transfected with same siRNA for 5-6hrs, and then were transferred to 35-mm wells. On day 4, the plasmid encoding the I-SceI endonuclease was cotransfected with the mCherry-expressing plasmid to induce DSBs. The cells were examined by flow cytometry on day 7, and the ratio of GFP to mCherry was used as a measure of HR or Alt-NHEJ efficiency. The C-NHEJ assay utilizes quantitative real-time PCR and was carried out as described (Hu and Parvin, 2014, Zhuang, et al., 2009) in 293 cells with modification. The transfection procedure was as described above. For plasmid add-back in the rescue assays, the transfection procedure was the same except that at the second siRNA transfection, the blank control plasmid, plasmids encoding Flag-TRF2 and corresponding mutants were co-transfected into the cells using Lipofectamine 3000 (Thermo Fisher Scientific). The data represent the means ± the s.d. of three separate experiments. P values were calculated by two-tailed t-test.

## Acknowledgments

We thank Dr. Jeff Parvin for kindly providing DSB repair assay cell lines, Dr. Eros Lazzerini Denchi for TRF mutant plasmids, and Dr. Feng Qiao for helpful discussion and advice.

## Funding

This work was supported by the National Science Foundation (MCB-1615701 to K.Y.) and AFOSR Grant # FA9550-08-1-0384 and gifts from the Beckman Laser Institute Foundation, the Hoag Family Foundation, and the David and Lucile Packard Foundation (M.W.B.), and by the Canadian Institutes of Health Research (MOP-86620 to X.Z.).

---

**Supplemental Figure S1.**
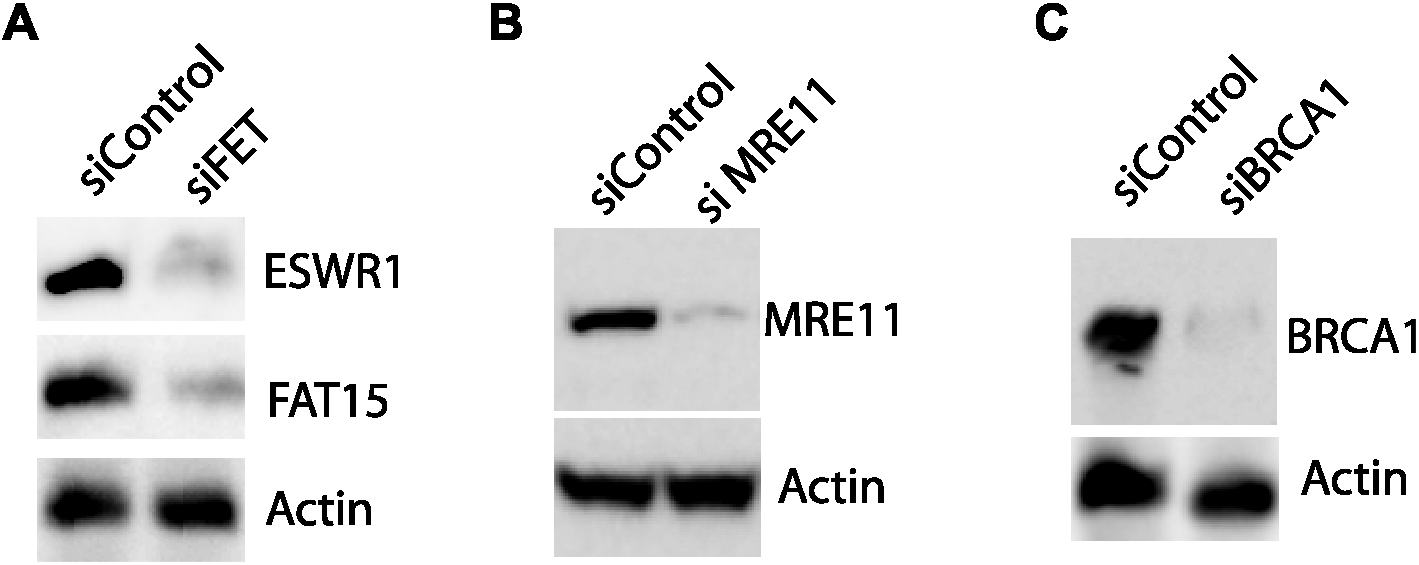
Co-depletion of ESWR1 and FET by siRNA specific for FET is confirmed by western blot (**A**). Actin serves as a loading control. Similarly, depletion of Mre11 (**B**) and BRCA1 (**C**) by corresponding siRNAs were confirmed.

**Supplemental Figure S2.**
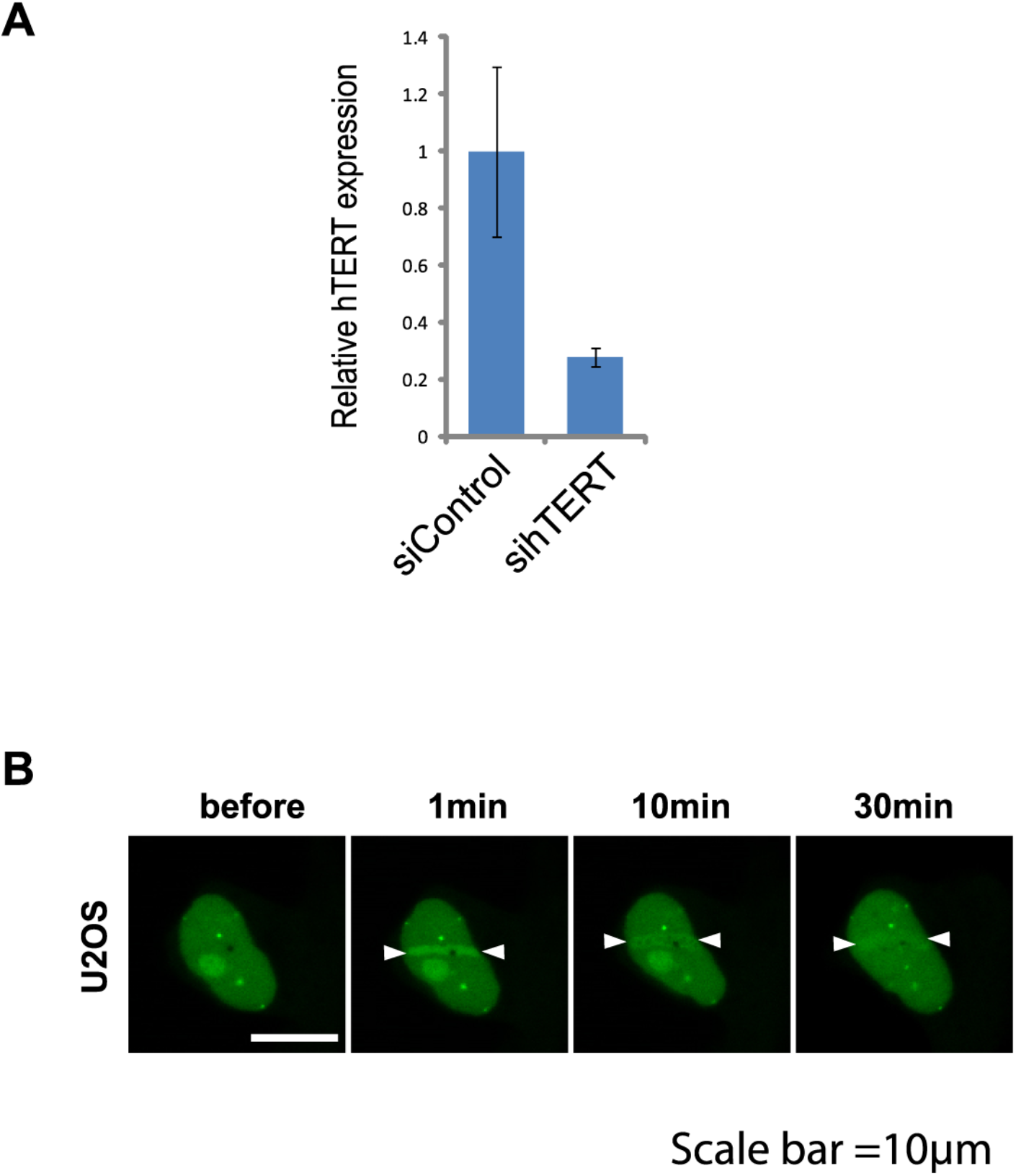
**A.** RT qPCR analysis of hTERT depletion in HeLa cells. The mRNA level of hTERT was normalized to GAPDH. hTERT qPCR primers used are 5’-CGGAAGAGTGTCTGGAGCAA-3’ (forward) and 5’-GGATGAAGCGGAGTCTGGA-3’(reverse) (Liu et al., 2013). **B.** An example of the time course analysis of GFP-TRF2 recruitment to the laser-induced damage sites in U2OS cells. Scale bar=10μm.

**Supplemental Figure S3.**
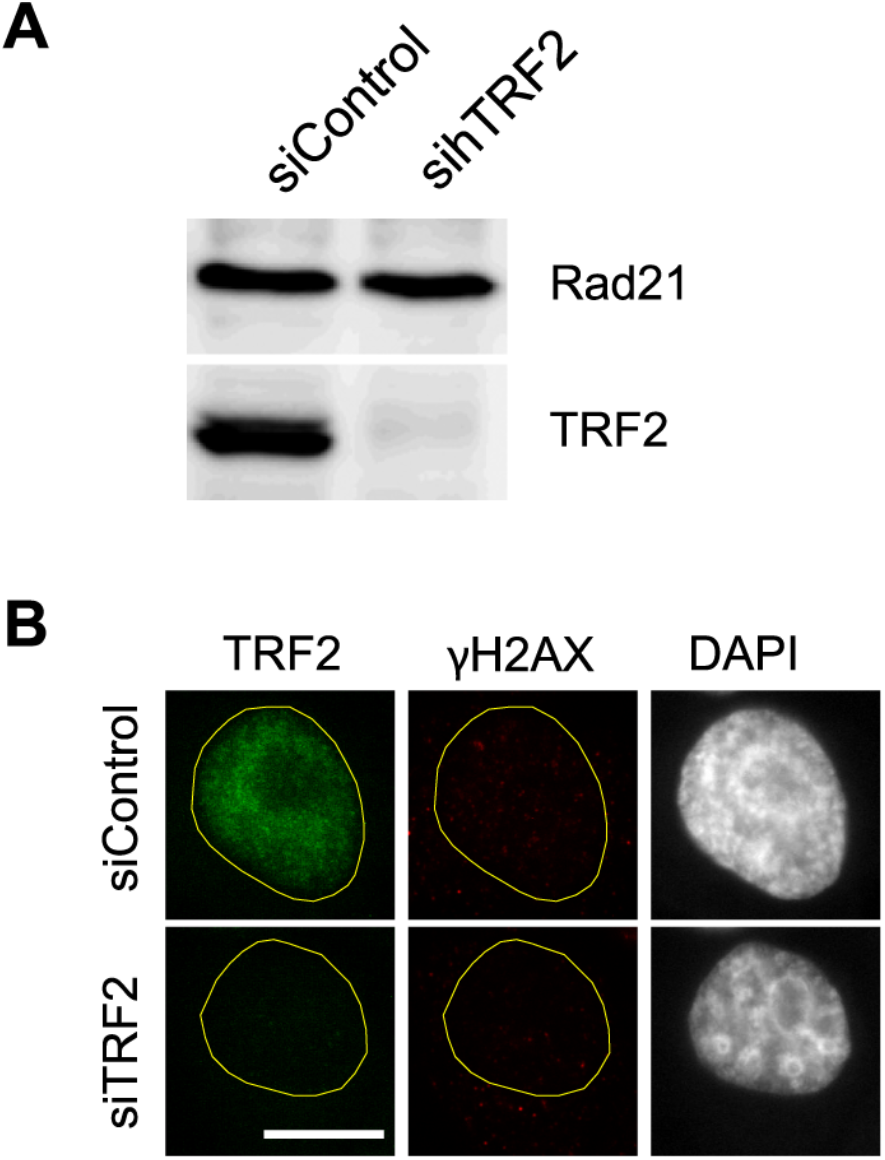
**A.** Analysis of TRF2 depletion in HeLa cells after siRNA transfection by Western blot. Total cell lysates from control or TRF2 siRNA-transfected cells were subjected to western blot analysis using anti-TRF2 antibody. Rad21 was used as a loading control. **B.** Telomere dysfunction-induced foci (TIF) analysis. Cells were treated with control or TRF2 siRNA and were fixed at 48hrs after 2nd siRNA transfection. Cells were stained with antibodies specific for TRF2 and γH2AX, and DAPI as indicated at the top. TRF2 depleted cells didn’t show any increase of γH2AX foci indicative of TIF.

